# Optogenetic control of gut bacterial metabolism to promote longevity

**DOI:** 10.1101/2020.02.25.964866

**Authors:** Lucas A. Hartsough, Matthew V. Kotlajich, Bing Han, Chih-Chun J. Lin, Lauren Gambill, Meng C. Wang, Jeffrey J. Tabor

**Author notes:** corresponding authorsemail.

## Abstract

Gut microbial metabolism is associated with host longevity. However, because it requires direct manipulation of microbial metabolism *in situ*, establishing a causal link between these two processes remains challenging. We demonstrate an optogenetic method to control gene expression and metabolite production from bacteria residing in the host gut. We genetically engineer an *Escherichia coli* strain that synthesizes and secretes colanic acid (CA) under the quantitative control of light. Using this optogenetically-controlled strain to induce CA production directly in the *Caenorhabditis elegans* gut, we reveal the local effect of CA in protecting intestinal mitochondria from stress-induced hyper-fragmentation. We also exploit different intensities of light to determine that the lifespan-extending effect of CA is positively correlated with its levels produced from bacteria. Our results show that optogenetic control offers a rapid, reversible and quantitative way to fine-tune gut bacterial metabolism and uncover its local and systemic effects on host health and aging.

## Introduction

Microbiome studies have identified correlations between bacteria and host aging (Kundu et al., 2017; O’Toole and Jeffery, 2015). For example, 16S rRNA and metagenomic DNA sequencing are used to associate the presence or abundance of specific bacteria to human centenarians (Biagi et al., 2016; Claesson et al., 2012, 2010). However, given the complexity and heterogeneity of the human gut environment, these approaches are unable to elucidate how a specific microbial species contributes to longevity. The nematode *Caenorhabditis elegans* has a short and easily-measured lifespan, features that have revolutionized our understanding of the molecular genetics of aging and longevity (Kenyon, 2010). Studies using *C. elegans* also provide mechanistic insight into the association between bacterial species and host longevity (Gusarov et al., 2013; Kim, 2013). Importantly, recent studies have revealed that bacterial metabolism can produce specific products to directly influence the aging process in the host *C. elegans* or modulate the effects of environmental cues on *C. elegans* lifespan (Cabreiro et al., 2013; Pryor et al., 2019; Virk et al., 2016). These findings highlight the significance of bacterial metabolism in regulating host physiology during the aging process and have inspired interest in directly manipulating bacterial metabolism *in situ* in the host gastrointestinal (GI) tract.

In several recent studies, researchers have administered antibiotic- or carbohydrate-based small molecule inducers to modulate gene expression from gut bacteria (Kotula et al., 2014; Lim et al., 2017; Mimee et al., 2015). While this approach has enabled the *in situ* analysis of a gut bacterial-host interaction (Lim et al., 2017), chemical effectors may have unwanted side-effects on host or microbial physiology and subject to slow and poorly-controlled transport and degradation processes that ultimately limit their precision.

Optogenetics combines light and genetically-engineered photoreceptors to achieve unrivaled control of biological processes (Olson and Tabor, 2014). Previously, we and others have engineered bacterial photoreceptors that activate or repress gene expression in response to specific wavelengths of light (Levskaya et al., 2005; Li et al., 2020; Ohlendorf et al., 2012; Ong and Tabor, 2018; Ong et al., 2017; Ramakrishnan and Tabor, 2016; Ryu and Gomelsky, 2014; Schmidl et al., 2014). These photoreceptors have been used to achieve precise quantitative (Olson et al., 2017, 2014), temporal (Chait et al., 2017; Milias-Argeitis et al., 2016; Olson et al., 2017, 2014), and spatial (Chait et al., 2017; Levskaya et al., 2005; Ohlendorf et al., 2012; Tabor et al., 2010) control of bacterial gene expression in culture conditions. They have also been used to characterize and control transcriptional regulatory circuits (Chait et al., 2017; Olson et al., 2014; Tabor et al., 2009) and bacterial metabolic pathways (Fernandez-Rodriguez et al., 2017; Tandar et al., 2019) *in vitro*. Here, we hypothesized that optogenetic control of bacterial gene expression might provide a new way to manipulate bacterial metabolism *in vivo* in the host GI tract, with high temporal and spatial precision and no unwanted side-effects.

We address this possibility using the *E. coli-C. elegans* interaction model, a testable system with known mechanistic links between bacterial metabolism and host longevity and with complete optical transparency. In particular, CA is an exopolysaccharide synthesized and secreted from *E. coli*, which can extend the lifespan of the host *C. elegans* through modulating mitochondrial dynamics (Han et al., 2017). We thus have genetically engineered an *E. coli* strain to put its biosynthesis of CA under a switchable control between green and red lights, and then utilized green light to induce CA production from this strain in the gut of the host *C. elegans.* We discovered that light-induced CA from bacteria residing in the host is sufficient to modulate mitochondrial dynamics and lifespan, which gives even more potent effects than dietary supplementation of CA. Furthermore, this optogenetic manipulation allowed us to investigate the local effect of CA on intestinal cells in a time-controlled manner and its systemic effect on organisms in a quantitative way, which are not possible with purified CA or CA-overproducing genetic mutants. This work paves the road for future application of bacterial optogenetics in understanding bacteria-host interaction with temporal, spatial and quantitative controls and minimal chemical interference.

## Results

To demonstrate optogenetic control over gut bacterial gene expression, we first engineered *E. coli* strain LH01, wherein our previous light-switchable two-component system CcaSR (Schmidl et al., 2014) controls expression of superfolder green fluorescent protein (*sfgfp*), and *mcherry* is expressed constitutively to facilitate identification of the bacteria (**Fig. 1a**, **Supplementary Fig. 1, Supplementary Tables 1-3**). In LH01, green light exposure switches CcaS to an active state in the presence of chromophore phycocyanobilin (PCB), wherein it phosphorylates the response regulator CcaR. Phosphorylated CcaR then activates transcription of *sfgfp* from the P_*cpcG2*-172_ output promoter. Red light reverts active CcaS to the inactive form, de-activating *sfgfp* expression (**Fig. 1a**).

**Figure 1.**
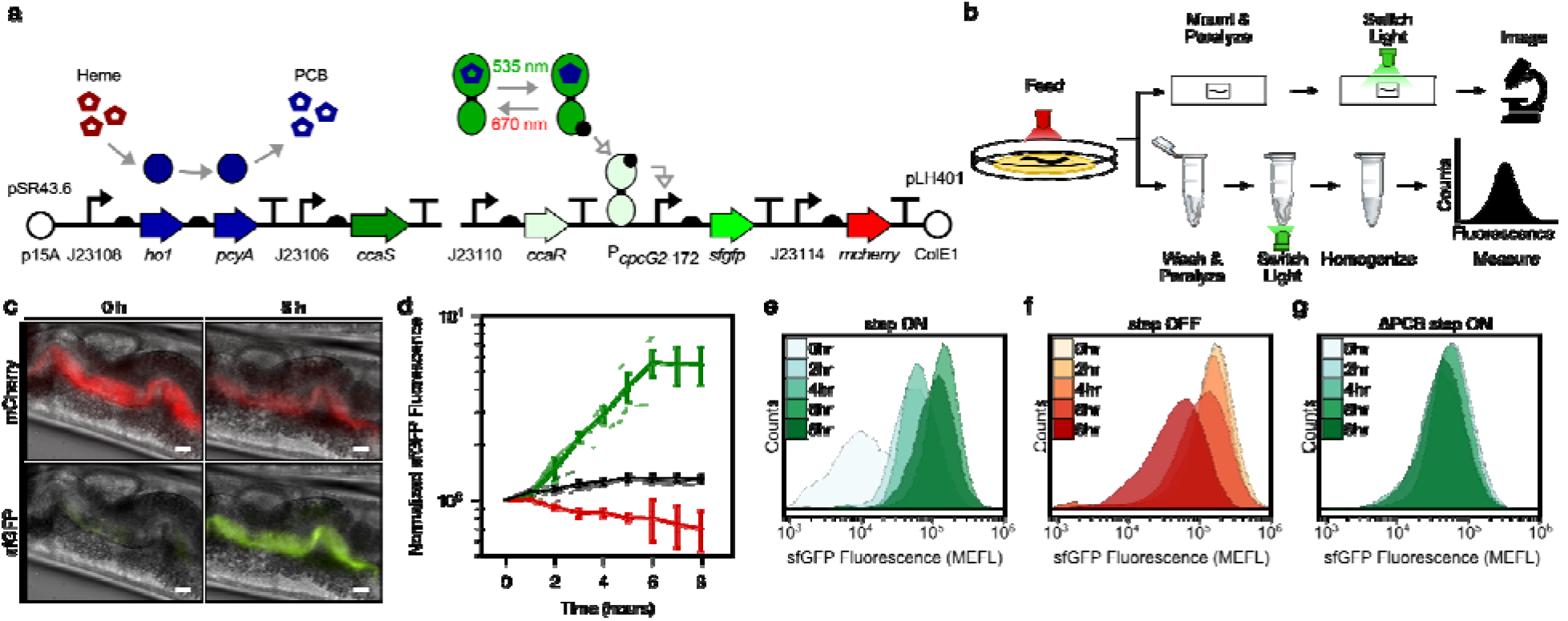
Optogenetic control of *C. elegans* gut bacterial gene expression. (a) Strain LH01. (b) Microscopy and cytometry workflows. (c) Fluorescence microscopy images 0 and 8 h after green light exposure in the step ON experiment. Scale bar: 10 μm. (d) Response dynamics in the step ON (green) and step OFF (red) microscopy experiments. Black: ΔPCB strain (step ON experiment). Individual-(light lines) and multi-worm average (dark lines) data are shown. *n* = 7, 4, 6 worms for green, red, black data sets (measured over 2, 3, 1 days, respectively). Error bars: SEM. (e-g) Flow cytometry histograms for response dynamics experiments. MEFL: molecules of equivalent fluorescein.

We then reared two groups of *C. elegans* from the larval to the adult stage on plates of LH01 under red or green light, respectively (**Fig. 1b**). Next, we washed away external bacteria, applied the paralyzing agent levamisole to prevent expulsion of gut contents, and transferred the worms to agar pads. Finally, we switched the light color from red to green, or green to red, and used epi-fluorescence microscopy to image the resulting changes in fluorescence in the gut lumen over time (**Fig. 1c**). In the red-to-green (step ON) experiment, we observed that sfGFP fluorescence in the worm gut lumen starts low, begins to increase within 2 hours, and reaches a saturated high level at 6 hours (**Fig. 1d**). In contrast, in the green-to-red (step OFF) experiment, sfGFP fluorescence begins high, and decreases exponentially between hours 1-7 (**Fig. 1d**). This light response is abolished when the PCB biosynthetic operon is removed (ΔPCB) (**Fig. 1d**), demonstrating that sfGFP levels are specifically controlled by CcaSR.

Next, we used flow cytometry to achieve high-throughput single-cell resolution analysis of this *in situ* bacterial light response. Specifically, we reared worms in red and green light as before, but then washed, paralyzed, and placed them into microtubes prior to light switching (**Fig. 1b**). At several time points over the course of 8 hours, we homogenized the animals, harvested the gut contents, and sorted bacterial cells and measured fluorescence via cytometry. This experiment revealed that bacteria in the host gut remain intact (**Supplementary Fig. 2**) and respond to light in a unimodal fashion (**Fig. 1e-g**). Furthermore, the temporal dynamics of the gene expression response and dependence on PCB recapitulate our microscopy results (**Supplementary Fig. 3)**. We also confirmed that residual bacteria on the exterior of worms do not contribute to the flow cytometry measurements (**Supplementary Fig. 4**). Together, these experiments demonstrate that we can use optogenetics to rapidly, reversibly induce gene expression of *E. coli* residing in the *C. elegans* gut.

Next, we sought to utilize our optogenetic method to modulate the production of specific metabolites in bacteria residing in the gut of live hosts. It is well-known that bacterial genes involved in the same metabolic process are often clustered into operons and co-regulated at the transcriptional level. We took advantage of this coordinated mode of regulation and chose the *cps* operon and its transcription activator RcsA for testing optogenetic control of bacterial metabolism. The *cps* operon in *E. coli* consists of 19 genes that encode enzymes required for the biosynthesis and secretion of CA (Torres-Cabassa and Gottesman, 1987), and CA-overproducing bacterial mutants Δ*lon* and Δ*hns* promote longevity in the host *C. elegans* (Han et al., 2017). To place CA biosynthesis under optogenetic control, we engineered an *E. coli* strain (MVK29) lacking genomic *rcsA* and expressing a heterologous copy of *rcsA* under the control of CcaSR (**Fig. 2a**, **Supplementary Tables 1-3)**. We first examined whether MVK29 could respond to green light and induce CA production and secretion. To this end, we grew the strain in batch culture under red or green light and quantified supernatant CA levels. In red light, MVK29 secretes CA to concentrations below the limit of detection of the assay, similar to the Δ*rcsA* mutant (**Fig. 2b**). Green light, on the other hand, induces MVK29 to secrete high levels of CA, and removal of the PCB biosynthetic operon abolishes this response (**Fig. 2b**). Moreover, mutation of the CcaS catalytic histidine to a non-functional alanine (H534A), or the CcaR phosphorylation site from an aspartic acid to a non-functional asparagine (D51N) abolishes detectable CA production (**Fig. 2b**). Importantly, the level of secreted CA increases sigmoidally with green light intensity, similar to the response of CcaSR itself (**Fig. 2c**) (Schmidl et al., 2014). We conclude that we have placed CA production under the control of the CcaSR system, and that we can use light to tune the production of bacterial metabolites.

**Figure 2.**
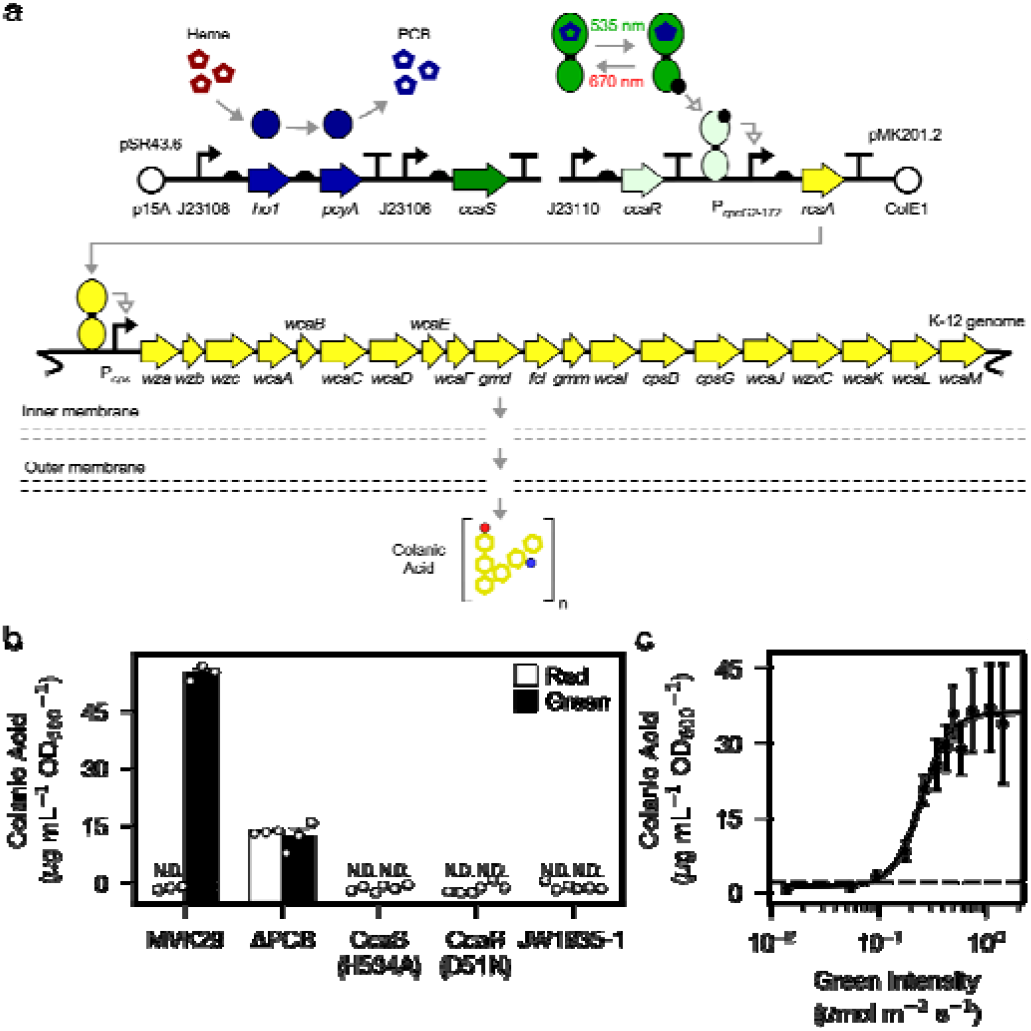
Optogenetic control of colanic acid biosynthesis. (a) Strain MVK29. (b) CA secretion levels for MVK29 and control strains exposed to red and green light. JW1935-1 is the *E. coli rcsA* background strain used in this study. N.D.: below assay limit of detection. (c) Green light intensity versus CA secretion level for MVK29. Data points represent 3 biological replicates collected on a single day. Dashed line: limit of detection. Error bars indicate standard deviation of the three biological replicates.

We then used this approach to study a gut bacterial metabolite-host interaction pathway *in vivo*. We first reared worms expressing mitochondrially-localized GFP (mito-GFP) (**Supplementary Table 3**) on MVK29 with red light. We then continued this red-light exposure for one group, and switched a second to green, for an additional 6 hours, and immediately imaged intestinal cell mitochondrial morphology using confocal microscopy (**Fig. 3b**). We found that mitochondrial fragmentation increases in worms exposed to the bacteria with light-induced CA secretion (**Fig. 3c**). This result recapitulates the phenotype observed in worms supplemented with purified CA (Han et al., 2017).

**Figure 3.**
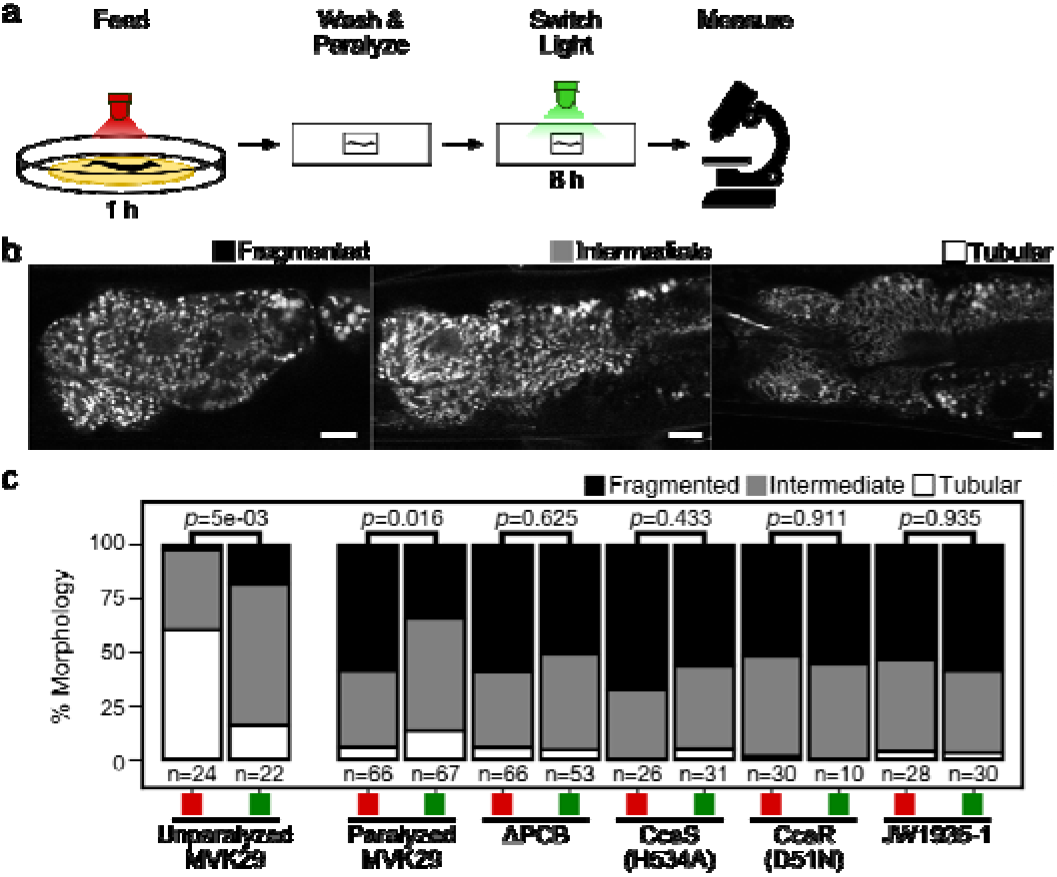
Light-regulated CA secretion modulates *C. elegans* mitochondrial dynamics. (a) Schematic of experiment for activating CA biosynthesis *in situ*. (b) Representative images of the mitochondrial network of anterior intestinal cells immediately distal to the pharynx are scored as fragmented, intermediate, or tubular, as previously(Han et al., 2017). Scale bars: 10 μm. (c) Mitochondrial fragmentation profiles of un-paralyzed worms fed MVK29 while exposed to red or green light for 6 h. (d) Fragmentation profiles for worms fed the indicated strain, then paralyzed for 6 h while exposed to red or green light. The number of worms included in each condition is indicated below each bar. The Chi-Squared Test of Homogeneity was used to calculate *p*-values between conditions.

Next, we induced CA production directly from bacteria residing within the host gut. To this end, we paralyzed the worms carrying mito-GFP, split them into two groups, and treated one with red and the second with green for 6 hours (**Fig. 3a**). We then analyzed mitochondrial morphology in intestinal cells using confocal microscopy. We found that the 6-hour levamisole treatment does not kill worms but leads to mitochondrial hyper-fragmentation (**Fig. 3d**), which might be due to the inhibitory effect of levamisole on mitochondrial NADH-oxidizing enzymes (Köhler and Bachmann, 1978). This stress-induced effect resembles mitochondrial decay related to aging and age-related neurodegenerative diseases (Cho et al., 2009; Exner et al., 2007; Sebastián et al., 2017). Interestingly, we found that green light exposure counteracts this hyper-fragmentation in paralyzed worms bearing MVK29 in the gut (**Fig. 3d**). Importantly, we found no such effects in worms bearing the ΔPCB, CcaS(H534A), CcaR(D51N) or *ΔrcsA* mutant strains (**Fig. 3d**), suggesting that this protective effective is a result of light-induced CA overproduction in the gut. These results not only show that optogenetics can be utilized to induce CA secretion from gut-borne *E. coli in vivo*, but also reveal a local protective effect of CA on intestinal cells.

Finally, we took the advantage of the quantitative control afforded by optogenetics to investigate how the lifespan-extending effect of CA relates to CA levels. Beginning at the day-1 adult stage, we exposed worms bearing MVK29 to red, or two green light intensities resulting in intermediate or high CA secretion, and measured their lifespans. We found that the lifespan extension increases proportionally with green-light intensity (**Fig. 4a**), revealing the pro-longevity effect of CA is dose dependent. We also noticed that the extent of lifespan extension by light is much stronger than that caused by dietary supplementation of purified CA (Han et al., 2017), suggesting the high efficacy of optogenetic induction to modulate host physiology. As a control, we repeated the experiment with worms bearing the CA-overproducing Δ*lon* mutant and showed that the lifespan extension caused by Δ*lon* is independent of light exposure (**Fig. 4b**). The extent of lifespan extension is similar to the MVK29 intermediate green light condition, but less than the MVK29 high green light condition (**Fig. 4a, b**), suggesting that MVK29 is capable of producing higher levels of CA than Δ*lon*. In addition, the lifespan of worms bearing the Δ*rcsA* mutant is also light-independent and is comparable to that of MVK29 worms under red light (**Fig. 4a, c**). These results suggest that optogenetic control is sufficient to induce bacterial production of pro-longevity compounds and improve host health, and can exert stronger beneficial effects than administration of a bacterial mutant or supplementing purified compounds. Importantly, unlike the traditional approach of introducing a bacterial mutant, optogenetic control of bacterial metabolism can modulate a host-level phenotype in a quantitative manner.

**Figure 4.**
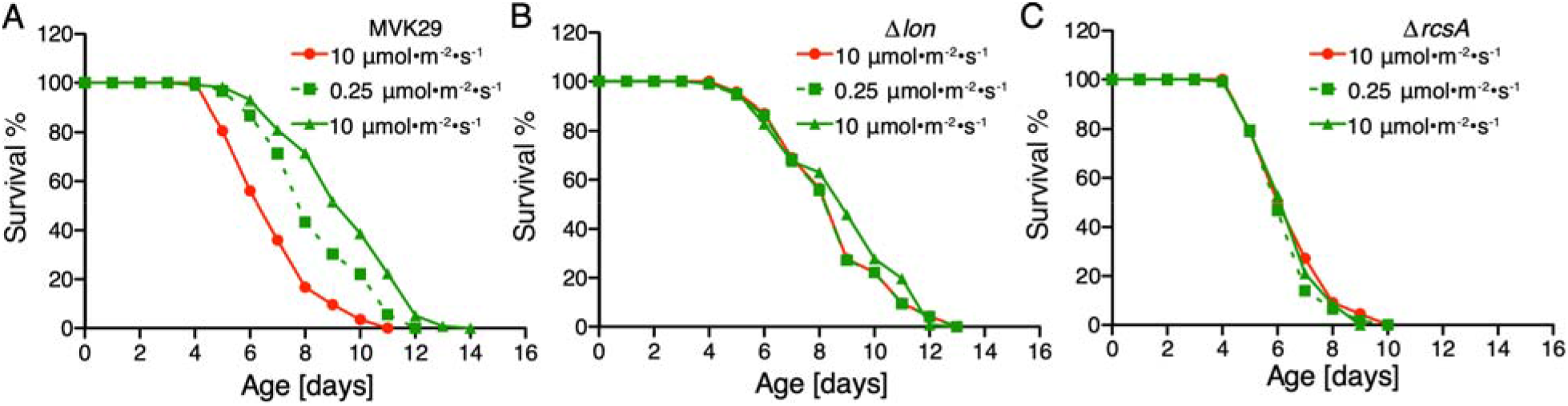
Optogenetically-regulated CA biosynthesis extends worm lifespan. (a) When exposed to green light, worms grown on MVK29 live longer than those exposed to red light, and the magnitude of lifespan extension is proportional to green light intensity (*p<0.0001* green vs. red, log-rank test). (b-c) The lifespans of worms grown on the Δ*lon* (b) or the Δ*rcsA* (c) controls are not affected by light condition (*p>0.1* green vs. red, log-rank test).

## Discussion

Our method has broad applications for studying microbe-host interactions *in situ*. For example, we have identified about two dozen additional *E. coli* genes that are unrelated to CA biosynthesis and that enhance worm longevity when knocked out (Han et al., 2017), though the mechanisms by which they act remain largely unclear. By using light to induce their expression in the gut and measuring acute host responses such as changes in mitochondrial dynamics, the role of these genes in gut microbe-host interactions could be further explored. In another example, the quorum-sensing peptide CSF and nitric oxide, both of which are produced by *Bacillus subtilis* during biofilm formation, have been found to extend worm lifespan through downregulation of the insulin-like signaling pathway (Donato et al., 2017). We have recently ported CcaSR into *B. subtilis* and demonstrated that it enables rapid and precise control of gene expression dynamics (Castillo-Hair et al., 2019). The method we report here should enable *in situ* studies of how gene expression and metabolite production from this important Gram-positive model bacterium impact longevity as well.

Multiple photoreceptors could also be combined to study more complex microbe-host interaction pathways. Specifically, we and others have co-expressed CcaSR with independently-controllable blue/dark and red/far-red reversible light sensors in order to achieve simultaneous and independent control of the expression of up to three genes in the same bacterial cell (Fernandez-Rodriguez et al., 2017; Olson et al., 2017; Tabor et al., 2010). Such optogenetic multiplexing could be performed *in situ* and used to study potential synergistic, antagonistic, or other higher-order effects of multiple bacterial genes or pathways. A large number of eukaryotic photoreceptors have also been developed, enabling optical control of many cell- and neurobiological processes (Deisseroth, 2015; Gautier et al., 2014; Goglia and Toettcher, 2018; Leopold et al., 2018). Bacterial and eukaryotic photoreceptors could be combined to enable simultaneous optical manipulation of bacterial and host pathways in order to interrogate whether or how they interact. Optogenetics could also be used to manipulate bacterial and/or host pathways at specific locations within the gut to examine location- or tissue-dependent phenomena.

Finally, our method could be extended to other bacteria or hosts. In particular, it should be possible to port CcaSR or other bacterial photoreceptors into native *C. elegans* symbionts (Zhang et al., 2017) or pathogens(Couillault and Ewbank, 2002). Because these strains stably colonize the host, the use of these bacteria could eliminate the need for paralysis, and facilitate longer-term experiments. It is likely that light can also be used to control gut bacterial gene expression in other model hosts such as flies, zebrafish, or mammals. Red-shifted wavelengths and corresponding optogenetic tools(Ong et al., 2017; Ryu and Gomelsky, 2014) may prove superior for less optically transparent or larger animals. Overall, by enabling precision control of bacterial gene expression and metabolism *in situ*, we believe that optogenetics will greatly improve our understanding of a wide range of microbe-host interactions.

## Acknowledgements

The LED array used to illuminate *C. elegans* plates was designed by Brian Landry & Sebastián Castillo-Hair. The mounting hardware for the microscope LED array was designed by Ravi Sheth. We thank Ravi Sheth for discussions during early stages of the project. We thank Dr. Joel Moake for the use of his cytometer. This work was supported by the John S. Dunn Foundation (J.J.T. and M.C.W.) and US National Institutes of Health, 1R21NS099870-01 (J.J.T.), DP1DK113644 (M.C.W.), R01AT009050 (M.C.W.). LAH was supported by a NASA Office of the Chief Technologist Space Technology Research Fellowship (NSTRF NNX11AN39H).

## Author Contributions

JJT and MW conceived of the study. LAH and MVK designed experiments. MVK and LAH constructed plasmids and strains. LAH, MVK, BH, CJL, LG, and MW performed experiments. CJL and LG scored single-blinded mitochondrial confocal micrographs. LAH, MVK, and MW analyzed and interpreted results. LAH, MW, and JJT wrote the manuscript.

## Methods

### *E. coli* plasmids, strains, and media

Plasmids used in this study are described in **Supplementary Table 1**. Genbank accession numbers are given in **Supplementary Table 2**. All plasmids constructed in this study were assembled via Golden Gate cloning(Engler et al., 2009). Primers were ordered from IDT (Coralville, IA). Assembled plasmids were transformed into *E. coli* NEB10β (New England Biolabs) for amplification and screening. All plasmid sequences were confirmed by Sanger sequencing (Genewiz; S. Plainfield, NJ). To construct pLH401 and pLH405, pSR58.6(Schmidl et al., 2014) was modified by inserting an *mCherry* expression cassette composed of a constitutive promoter (J23114; http://parts.igem.org/Promoters/Catalog/Anderson), RBS (BBa_B0034; http://parts.igem.org/Part:BBa_B0034), *mCherry*, and a synthetic transcriptional terminator (L3S1P52 (Chen et al., 2013)). To construct pLH405, pLH401 was further modified by exchanging the superfolder GFP gene (*sfgfp*) for *gfpmut3\**. pMVK201.2 was built by modifying pSR58.6 to control expression of *rcsA*.

All *E. coli* strains are described in **Supplementary Table 3.** *ΔrcsA* (JW1935-1) was obtained from the Coli Genetic Stock Center. *Δlon* (JW0429-1) was obtained from the Keio *E. coli* knockout library(Baba et al., 2006), a gift from the Herman lab. All *E. coli* strains were maintained in LB media supplemented with appropriate antibiotics (chloramphenicol 34 μg/mL, spectinomycin 100 μg/mL, kanamycin 100 μg/mL) in a shaking incubator at 37°C and 250 rpm unless otherwise noted.

### *C. elegans* strains and media

All *C. elegans* strains (**Supplementary Table 3**) were provided by the Caenorhabditis Genetics Center (University of Minnesota), which is funded by the NIH office of Research Infrastructure Programs (P40 OD010440). Worms were grown at 20°C on 1.7% NGM-agar plates in 60 mm Petri dishes inoculated with a lawn of *E. coli* (CGSC str. BW28357), as described in the CGC WormBook (wormbook.org), unless otherwise specified. The common strain *E. coli* OP50 was not used for worm feeding, as it produces CA during normal growth(Han et al., 2017). M9 buffer for *C. elegans* (abbreviated M9Ce to distinguish from *E. coli* M9 media) was composed of 3 g KH_2_PO_4_, 6 g Na_2_HPO_4_, 5 g NaCl, 1 mL 1 M MgSO_4_, H_2_O to 1 L, and sterilized by autoclaving (wormbook.org).

### Optogenetic control of CA production

3 mL starter cultures of appropriate *E. coli* strains were inoculated from a −80°C freezer and grown 12 h at 37°C. These starters were diluted to OD_600_ = 1×10^−2^ in M9 minimal media (1x M9 salts, 0.4% w/v glucose, 0.2% w/v casamino acids, 2 mM MgSO_4_, 100 μM CaCl_2_) supplemented with appropriate antibiotics. The M9/cell mixtures were then distributed into 3 mL aliquots in 15 mL clear polystyrene culture tubes and grown at 37°C in a shaking incubator at 250 rpm while illuminated with the appropriate light wavelength and intensity, using the Light Tube Array (LTA) device (Gerhardt et al., 2016). After 22 h, cultures were removed and iced to halt growth and the OD_600_ was measured. Culture samples were collected for CA quantification.

### CA quantification

We adapted a previous CA quantification protocol(DISCHE, 1947; DISCHE and SHETTLES, 1948) that takes advantage of the fact that it is the only exopolysaccharide produced in our *E. coli* strains that incorporates fucose. In particular, we quantified the amount of fucose in cell-derived exopolysaccharides (EPS), and used that value as a proxy CA levels. First, EPS was liberated from cells by boiling 2 mL of culture for 15 min. in a 15 mL conical tube. The sample was then centrifuged in 1.5 mL Eppendorf tubes for 15 min. at 21,000 × g. Then, 0.7 mL of supernatant was dialyzed against water for at least 12 h using Pur-A-Lyzer Midi 3500 dialysis mini-tubes (Sigma-Aldrich, PURD35100-1KT) to remove monomeric fucose from the sample.

Fucose monomers were then liberated from the EPS polymers by hydrolyzing 0.2 mL of dialyzed media with 0.9 mL of H_2_SO_4_ solution (6:1 v/v acid:water). This mixture was boiled in a 15 mL conical for 20 min and then cooled to room temperature. The absorbance at 396 nm and 427 nm was measured. Next, 25 μL of 1 M L-cysteine HCl was added and mixed thoroughly by pipetting. The absorbance at 396 nm and 427 nm was measured again. Simultaneously, absorbance measurements of L-fucose standards pre- and post-L-cysteine addition were also recorded. Absorbance change, given by *D* in the formula below, were used to compare the L-fucose standard samples to the dialyzed culture samples and estimate the L-fucose concentration in the dialyzed product.

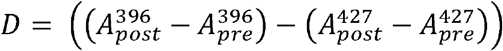

### Preparation of NGM-agar plates for worm feeding

3 mL *E. coli* starter cultures were inoculated from −80°C freezer stocks and grown for 12 h at 37°C. These starters were then diluted to OD_600_ = 1 × 10^−6^ in M9 minimal media supplemented with appropriate antibiotics. The M9/cell mixture was then distributed into 3mL aliquots in 15 mL clear polystyrene culture tubes and grown at 37°C in a shaking incubator at 250 rpm while illuminated with the appropriate light in the LTA. Once cultures reached OD_600_ = 0.1-0.4, tubes were iced for 10 min and subsequently concentrated to OD_600_ ~ 20 by centrifugation (4°C, 4000 rpm, 10 min) and resuspension in fresh M9 media. 400-600 μL of dense bacterial culture was then applied to sterile NGM-agar plates and allowed to dry in a dark room, or a room with green overhead safety lights if cultures were preconditioned in green light. Plates were wrapped in foil and refrigerated at 4°C for no more than 5 days until needed.

### Time-lapse microscopy

To obtain age-synchronized worm cultures, axenized *C. elegans* (strain *glo-1*) eggs were isolated and allowed to arrest in L1 by starvation in M9 buffer (distinct from M9 media: 3 g KH_2_PO_4_, 6 g Na_2_HPO_4_, 5 g NaCl, 1 mL 1M MgSO_4_, and water to 1 L, sterilized by autoclaving at 121°C for 20 min) at room temperature for 12-18 h. 10-100 larvae were transferred to a previously prepared NGM-agar plate containing a lawn of the appropriate bacterial strain. The plate was then placed in a 20°C incubator and illuminated with appropriate optogenetic light provided by a single LED positioned 1cm above the Petri dish. Adult worms were transferred to a fresh plate as necessary to maintain only a single generation.

Individual worms aged 1-3 days were removed from the dish and prepared for time-lapse epifluorescence imaging. A 1.5% agar pad was prepared using M9 buffer as previously described^45^, and punched into ½” circles with a hollow punch. A 4 μL droplet of 2 mM levamisole was deposited on a pad and 5 adult worms were transferred from the NGM plate to the droplet. An additional 4 μL of levamisole solution was added and the worms were gently washed to remove external bacteria. Worms were then transferred to a fresh pad with a 4 μL droplet of levamisole solution, which was allowed to dry, thereby co-localizing and aligning the worms on the pad. The pad was then inverted and placed into a 13 mm disposable microscopy petri dish with a #1.5 coverslip on the bottom (Cell E&G; Houston, TX). Another coverslip was placed on the top of the pad in the dish to curtail evaporation.

The dish was then placed on the stage of a Nikon Eclipse Ti-E inverted epifluorescence microscope (Nikon Instruments, Inc; Melville, NY). Complete paralysis was induced by incubating the dish at room temperature (~23°C) for 30 min. Meanwhile, worms were exposed appropriate preconditioning light supplied by a circular array of 8 LEDs (4 × 660 nm, 4 × 525 nm) mounted to the microscope condenser ring, about 2 cm above the Petri dish. Light was then switched from the preconditioning to the experimental wavelength, and worms were imaged periodically using 10x, 40x, and 60x objectives. For each time point, the LEDs were turned off and images acquired in the brightfield (DIC) and fluorescent channels. Afterwards, the LEDs were turned on again to maintain optogenetic control.

### Epifluorescence image analysis

All epifluorescence images were analyzed using the Nikon Elements software package (Nikon Instruments, Inc; Melville, NY). The mCherry signal was used as a marker for the gut lumen, and only cells in this region were included in the analysis. Image ROIs were created by thresholding the sfGFP signal to identify the boundaries of cell populations. Out of focus regions were eliminated from analysis. The average sfGFP pixel intensity inside the ROIs was calculated and recorded for each time point.

### Flow cytometry

1-3 day old *glo-1* worms were prepared for flow cytometry of the microbiome constituents by washing, using a protocol adapted from previous work(Portal-Celhay et al., 2012). Groups of 5 worms were washed 2x in a 5 μL droplet of lytic solution: *C. elegans* M9 buffer containing 2 mM levamisole, 1% Triton X-100, and 100 mg/mL ampicillin. The worms were then washed 2x in 5 μL droplets of M9 buffer containing 2 mM levamisole only. Finally, the worms were transferred to clear 0.5 mL Eppendorf tubes containing 50 μL of M9 buffer + 2 mM levamisole, ensuring that 5 worms were deposited in the liquid contained in each tube. Each tube was then exposed to light by placing it within one well of a 24-well plate (AWLS-303008, ArcticWhite LLC) atop a Light Plate Apparatus (LPA) containing green and red LEDs^47^ for 8 h at room temperature. In separate control experiments, we demonstrated that any stray bacteria that may escape the worms over this period, or which were inadvertently added to the 50 μL of M9 buffer, are incapable of responding to optogenetic light (**Supplementary Fig. 4**).

At the conclusion of the experiment, tubes were removed from the plate and immediately chilled in an ice slurry for 10 min in the dark. Worms were homogenized using an anodized steel probe sterilized between samples via 70% ethanol treatment and flame before being cooled.

Next, we used our previous antibiotic-based fluorescent protein maturation protocol(Olson et al., 2014) to allow unfolded proteins to mature while preventing the production of new protein. In particular, 250 μL PBS containing 500 mg/mL Rifampicin was added to the 50 μL homogenized worm samples and transferred to cytometry tubes. These tubes were incubated in a 37°C water bath for precisely 1 h, then transferred back to an ice slurry.

These samples were measured on a BD FACScan flow cytometer. For gating, an FSC/SSC polygon gate was first created using non-fluorescent bacteria grown *in vitro* at 37°C (**Supplementary Fig. 2**). Events outside this region were excluded as non-bacterial material. To isolate the engineered gut bacteria, only events with a high mCherry signal (FL3 > 1200 a.u., FL3 gain: 999) were included (**Supplementary Fig. 2**). Samples were measured until 20,000 events were recorded or the sample was exhausted.

### Flow cytometry data analysis

All flow cytometry data (FCS format) were analyzed using FlowCal(Castillo-Hair et al., 2016) and Python 2.7. We wrote a standard cytometry analysis workflow that truncated the initial and final 10 events to prevent cross-sample contamination, removed events from saturated detector bins at the ends of the detection range, and added 2D density gate on SSC/FSC retaining the densest 75% of events (**Supplementary Fig. 2a**). GFPmut3* fluorescence units were converted into standardized units of molecules of equivalent fluorescein (MEFL) using a fluorescent bead standard (Rainbow calibration standard; cat. no. RCP-30-20A, Spherotech, Inc.) as described previously(Castillo-Hair et al., 2016). Finally, to eliminate events associated with *C. elegans* autofluorescence (**Supplementary Fig. 2b**), any events in the region FL1 ≤ 1200 MEFL were discarded.

### Mitochondrial Fragmentation Assays

Synchronized L1 worms (strain *ges-1*) were applied to NGM agar plates containing bacterial strain BW25113 and allowed to develop until adulthood. This parental bacterial strain is used to allow all worms to develop at the same rate, avoiding any developmental/growth effects the experimental strains may exert on the worms. All experimental bacterial strains were preconditioned in red light with optogenetic light provided by a single LED positioned 1 cm above the Petri dish.

After 3-5 days (between days 1-3 of adulthood), worms allocated for the experiment were transferred to experimental strains for approximately 60-90 minutes to thoroughly inoculate the GI tract. In the case of the unparalyzed worms, red or green light was then applied for an additional 6 hours. For the paralyzed worm experiments, 1.5% low-melt agar pads were prepared as described above and placed on individual slides. About 15 adult worms were transferred from the experimental strain Petri dish to an agar pad containing 10 μL of *C. elegans* M9 buffer + 2 mM levamisole (M9Ce+Lev), where worms were gently washed before being transferred to a fresh pad also containing 10 μL of M9Ce+Lev. The majority of M9Ce+Lev on the pad was allowed to evaporate, which causes the worms to align longitudinally before a cover slip was applied. Slides were then exposed to either red or green light by placing them under a single LED positioned 1 cm above the Petri dish for 6 h. Afterward, the anterior intestinal cells were imaged using confocal microscopy (Olympus Fluoview 3000) in the brightfield and GFP channels.

### Confocal Microscopy Image Analysis

All confocal images for an experiment were manually cropped to display only the anterior intestinal cells of a single worm (in the GFP channel). These cropped images were then anonymized, randomized and the mitochondrial networks of each were blindly classified by two researchers independently as either tubular, fragmented, or intermediate. Tubular samples were marked by a high degree of network connectivity throughout. Fragmented samples were composed almost exclusively of isolated clusters of fluorescence with high circularity. Intermediate samples contained regions of both types. Scores were then de-randomized and aggregated. For each experimental strain, the red and green light conditions were compared for statistical significance using the chi-squared test of homogeneity.

### Lifespan experiments

3 mL starter cultures of Δ*lon*, Δ*rcsA* or MVK29 were inoculated from −80°C freezer stocks into LB supplemented with appropriate antibiotics and grown shaking for 12 h at 37 °C at 250 rpm. These cultures were diluted to OD_600_ = 1×10^−6^ in 27 mL M9 media supplemented with appropriate antibiotics. 1.5 mL of each M9/cell mixture was added to each of 18 wells on three 24-well plates and grown in 3 LPA devices under the appropriate light conditions at 37 °C and 250 rpm. Once cultures reached OD_600_ = 0.1-0.4, all tubes were iced for 10 min and subsequently concentrated 10 times by centrifugation (4°C, 4000 rpm, 10 min). Approximately 50 μL of this dense bacterial culture was then applied to sterile NGM-agar plates with no antibiotics and allowed to dry in a dark room. The plates were then illuminated with the appropriate light wavelength and intensity for 16 h at room temperature, and immediately used for lifespan assays.

During the longitudinal lifespan assay, exposure to white light is limited to the minimal level. To reach this goal, the *sqt-3(e2117)* temperature sensitive mutant (**Supplementary Table 3**) was used to perform longitudinal analyses at 25 °C, which avoids time-consuming animal transfer without interrupting normal reproduction. *sqt-3(e2117)* is a collagen mutant of *C. elegans* that reproduces normally but is embryonically lethal at 26 °C, and has been used previously in longitudinal studies(Han et al., 2017; Wang et al., 2014). Worms were age-synchronized by bleach-based egg isolation followed by starvation in M9 buffer at the L1 stage for 36 hours. Synchronized L1 worms were grown on BW25113 *E. coli* at 15 °C until the L4 stage, when worms were transferred to 24-well plates (~15 worms/well) with Δ*lon*, Δ*rcsA* or MVK29 (**Supplementary Table 3**). The plates were placed in LPA. The LPA LEDs were programmed to illuminate wells with constant red (10 μmol/m^2^/s), low-intensity green (0.25 μmol/m^2^/s), or high-intensity green light (10 μmol/m^2^/s). The apparatus was then transferred to a 26 °C incubator. The number of living worms remaining in each well was counted every day. Death was indicated by total cessation of movement in response to gentle mechanical stimulation. Statistical analyses were performed with SPSS (IBM Software) using Kaplan-Meier survival analysis and the log-rank test (**Supplementary Table 4**).

## Supplemental Information Titles and Legends

**Supplementary Figure 1. *In vitro* characterization of GFP reporter strains used in this study.** All optogenetic strains carry our previously published CcaSR v2.0 system, which is encoded on plasmids (a) pSR43.6 and (b) pSR58.6. (c) pLH401 (used for microscopy) and (d) pLH405 (used for cytometry) genetic device schematics. We replaced *sfgfp* with *gfpmut3\** in pLH405 as we hypothesized that the latter may be less stable and thus result in faster response dynamics. However, we observed no difference in dynamics. (e) Batch culture light responses of CcaSR v2.0 and all GFP reporter strains used in this study. GFP fluorescence was measured by flow cytometry. The dynamic range (ratio of GFP fluorescence in green versus red light) is shown above each data set. We note that CcaSR v2.0 exhibits 77-fold dynamic range in the reference strain BW29655 (Δ*envZ*, Δ*ompR*), which is similar to our previous measurement of this strain at 120-fold (Schmidl et al., 2014). The calculated fold-change is sensitive to fluctuations in the measured *E. coli* autofluorescence and red-light expression level, which likely explains the slight discrepancy. CcaSR v2.0 dynamic range increases slightly to 84.2-fold in Δ*rcsA* (JW1935-1), which is used throughout this work, due to higher sfGFP expression in green light. The mCherry cassette in pLH401 decreases dynamic range to 38.3-fold due to higher leaky sfGFP expression in red light. In worms, the fold-change in response to green light decreases further to 5.52 ± 2.4-fold (**Fig. 1d**). The use of *gfpmut3\** in pLH405 further decreases dynamic range to 13.5 ± 0.0-fold for reasons that are not clear, while in worms the fold-change is 8.63 ± 3.6-fold (**Fig. 1e**). Data represent the mean of three independent, autofluorescence-subtracted, biological replicates acquired on 3 separate days. Error bars: standard deviation.

**Supplementary Figure 2. Flow cytometry gating.** (a) Strain LH05 (**Supplementary Table 3**), which expresses only mCherry, was fed to worms, isolated, and analyzed by flow cytometry to quantify *E. coli* autofluorescence through the FL1 (GFP) channel. To eliminate events corresponding to cytometer noise and autofluorescence of homogenized worm samples, three gates were then applied: (1) a density gate for the most homogeneous 75% of samples in forward scatter (FSC) vs side-scatter (SSC) (the area within the bold black line in the plots in Column 1), (2) a threshold gate on the FL1 (GFP) channel (marked by a red dashed line; **Methods**), and (3) a threshold gate for events exhibiting high FL3 (mCherry, red dashed line; **Methods**). (b) Applying these gates to samples from the step ON experiment (**Fig. 1e**) reveals robust isolation of bacteria with a FSC/SSC profile consistent with our previous *in vitro* experiments^24,47^ and an increase in expression of GFPmut3* in response to green light. Ungated data are shown for comparison to gated data in all plots.

**Supplementary Figure 3. Gene expression dynamics from flow cytometry experiments.** (a) LH03 and (b) LH04 (ΔPCB) step ON results. (c) LH03 and (d) LH04 step OFF results. Each individual trajectory (faint lines) corresponds to a single biological replicate. A biological replicate comprises the homogenized contents of five worms collected on a single day. Each biological replicate on a given plot was run on a different day. Population medians taken across all trajectories are shown in bold (**Methods**). Data are composed of six and eight trajectories in the step ON and step OFF experiments, respectively. Error bars: SEM.

**Supplementary Figure 4. Bacteria on the exterior of worms do not contribute to the measured light response in flow cytometry experiments.** Worms were suspended in clear tubes containing M9Ce+Lev media and levamisole in the flow cytometry experiments. To demonstrate that bacterial cells outside the worm gut (e.g. on the exterior of the worm) do not respond to light in these conditions, samples of pre-conditioned bacteria from the NGM plates were suspended in the M9Ce+Lev or *E. coli* M9 media supplemented with casamino acids and glucose (M9Ec). These samples were then exposed to either green or red light for 8 h, then measured via flow cytometry after the fluorophore maturation protocol (**Methods**). As expected, the bacterial populations are only responsive to light when grown in M9Ec and when the PCB operon is intact (strain JW1935-1/pLH405/pSR43.6, aka LH03). We conclude that the responses shown in data in **Fig. 1e-f** and **Supplementary Fig. 3** are not due to bacteria outside the worm, nor to bacteria that escape the worm over the course of the experiment, as such cells do not respond to light in the experimental buffer (M9Ce). Data represent 7, 6, 3, and 5 replicates (left); and 8, 3, 8, and 3 replicates (right) over 8 and 10 days, respectively; error bars: SEM. Individual data points for each condition are overlaid as white markers.

**Supplementary Table 1. Plasmids used in this study**

**Supplementary Table 2. Genbank Accession Numbers**

**Supplementary Table 3. Bacterial and worm strains used in this study**

**Supplementary Table 4. Statistical analysis of worm lifespan experiments**

