## Supplementary Figures for "Optogenetic control of gut bacterial metabolism to promote longevity"

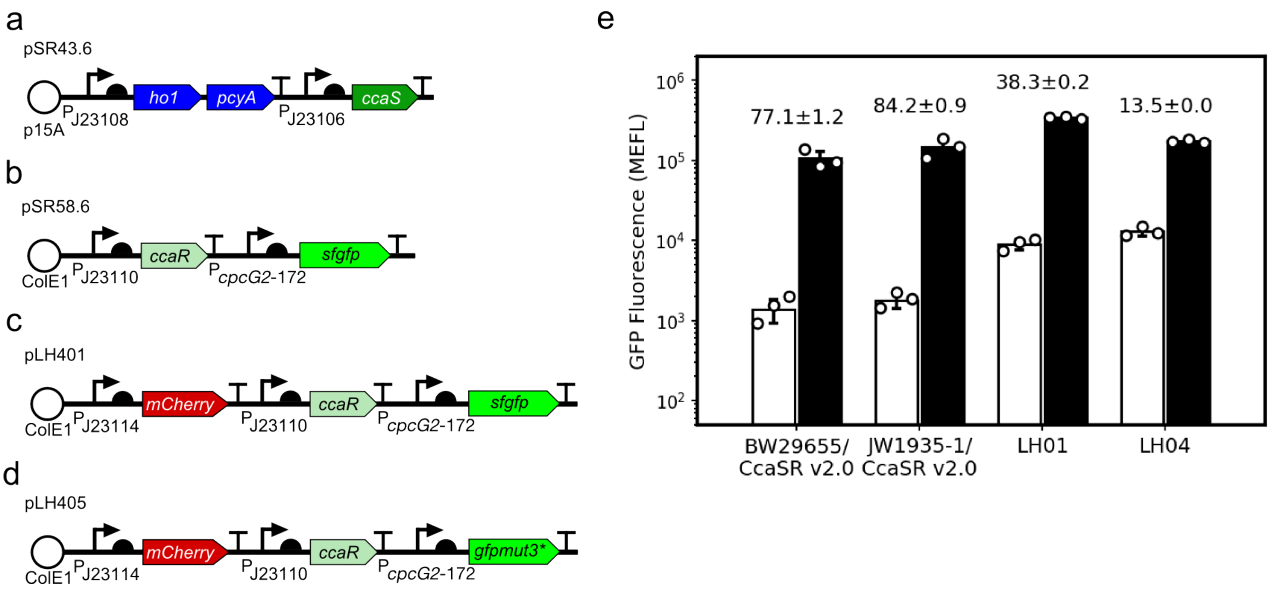


**Supplementary Figure 1. *In vitro* characterization of GFP reporter strains used in this study.** All optogenetic strains carry our previously published CcaSR v2.0 system, which is encoded on plasmids (a) pSR43.6 and (b) pSR58.6. (c) pLH401 (used for microscopy) and (d) pLH405 (used for cytometry) genetic device schematics. We replaced *sfgfp* with *gfpmut3** in pLH405 as we hypothesized that the latter may be less stable and thus result in faster response dynamics. However, we observed no difference in dynamics. (e) Batch culture light responses of CcaSR v2.0 and all GFP reporter strains used in this study. GFP fluorescence was measured by flow cytometry. The dynamic range (ratio of GFP fluorescence in green versus red light) is shown above each data set. We note that CcaSR v2.0 exhibits 77-fold dynamic range in the reference strain BW29655 (Δ*envZ*, Δ*ompR*), which is similar to our previous measurement of this strain at 120-fold[15]. The calculated fold-change is sensitive to fluctuations in the measured *E. coli* autofluorescence and red-light expression level, which likely explains the slight discrepancy. CcaSR v2.0 dynamic range increases slightly to 84.2-fold in Δ*rcsA* (JW1935-1), which is used throughout this work, due to higher sfGFP expression in green light. The mCherry cassette in pLH401 decreases dynamic range to 38.3-fold due to higher leaky sfGFP expression in red light. In worms, the fold-change in response to green light decreases further to 5.52 ± 2.4-fold (**Fig. 1d**). The use of *gfpmut3** in pLH405 further decreases dynamic range to 13.5 ± 0.0-fold for reasons that are not clear, while in worms the fold-change is 8.63 ± 3.6-fold (**Fig. 1e**). Data represent the mean of three independent, autofluorescence-subtracted, biological replicates acquired on 3 separate days. Error bars: standard deviation.

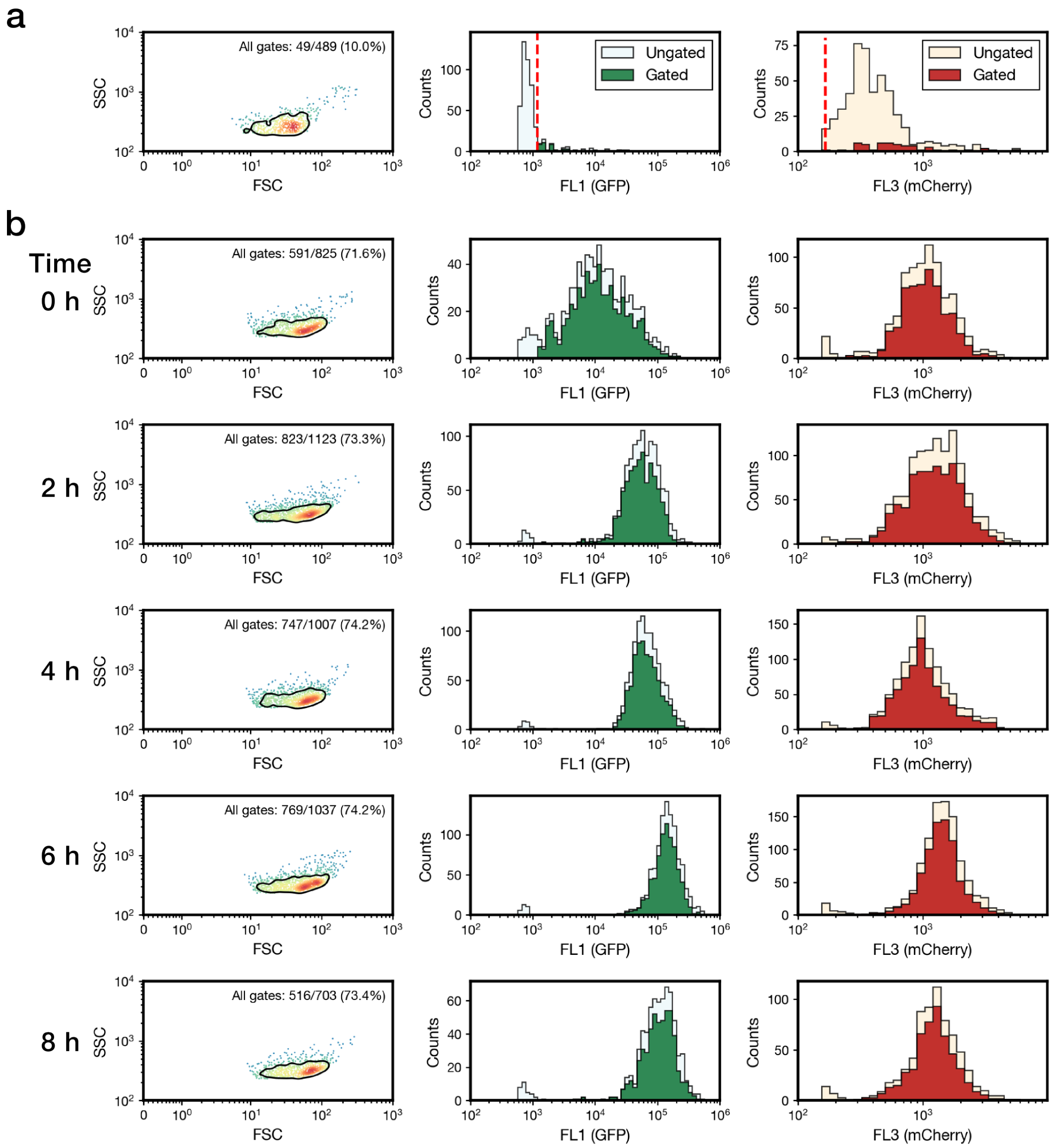


**Supplementary Figure 2. Flow cytometry gating.** (a) Strain LH05 (**Supplementary Table 3**), which expresses only mCherry, was fed to worms, isolated, and analyzed by flow cytometry to quantify *E. coli* autofluorescence through the FL1 (GFP) channel. To eliminate events corresponding to cytometer noise and autofluorescence of homogenized worm samples, three gates were then applied: (1) a density gate for the most homogeneous 75% of samples in forward scatter (FSC) vs side-scatter (SSC) (the area within the bold black line in the plots in Column 1), (2) a threshold gate on the FL1 (GFP) channel (marked by a red dashed line; **Methods**), and (3) a threshold gate for events exhibiting high FL3 (mCherry, red dashed line; **Methods**). (b) Applying these gates to samples from the step ON experiment (**Fig. 1e**) reveals robust isolation of bacteria with a FSC/SSC profile consistent with our previous *in vitro* experiments[24,47] and an increase in expression of GFPmut3* in response to green light. Ungated data are shown for comparison to gated data in all plots.

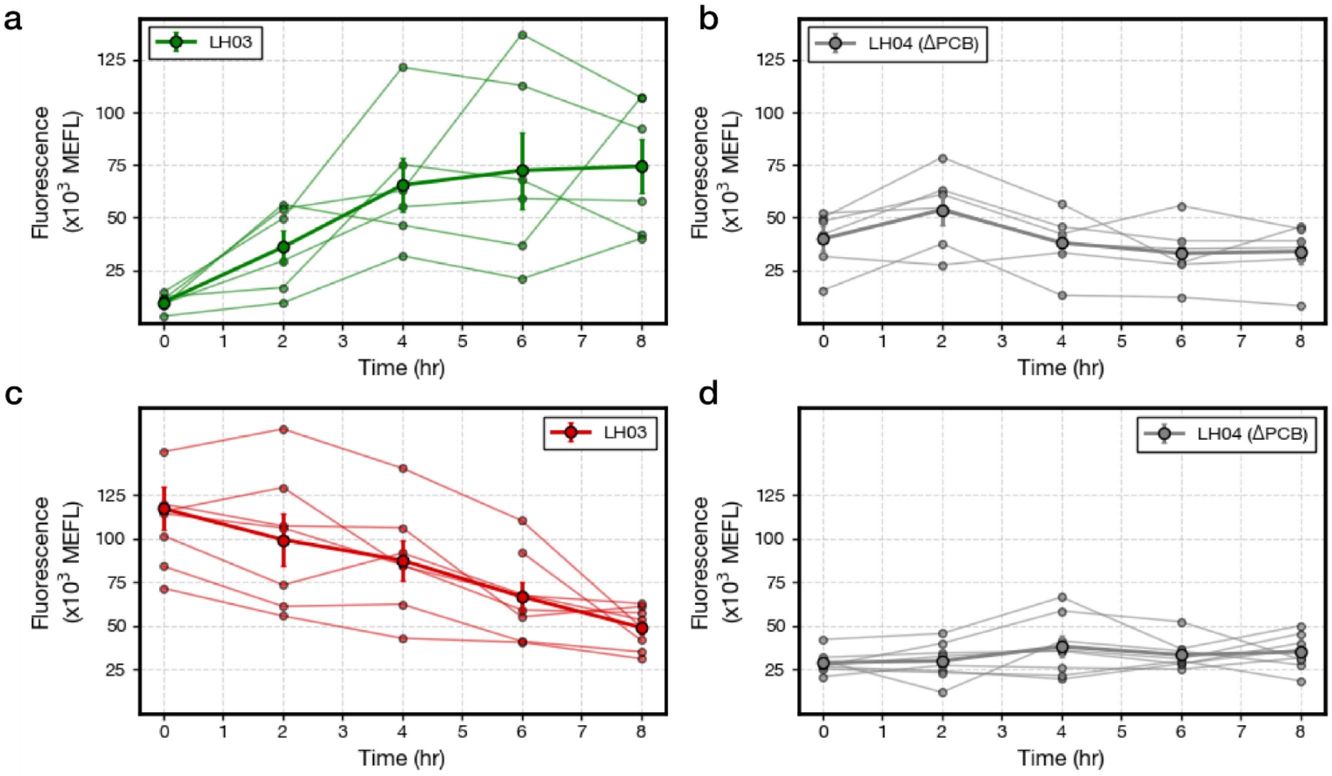


**Supplementary Figure 3. Gene expression dynamics from flow cytometry experiments.** (a) LH03 and (b) LH04 (ΔPCB) step ON results. (c) LH03 and (d) LH04 step OFF results. Each individual trajectory (faint lines) corresponds to a single biological replicate. A biological replicate comprises the homogenized contents of five worms collected on a single day. Each biological replicate on a given plot was run on a different day. Population medians taken across all trajectories are shown in bold (**Methods**). Data are composed of six and eight trajectories in the step ON and step OFF experiments, respectively. Error bars: SEM.

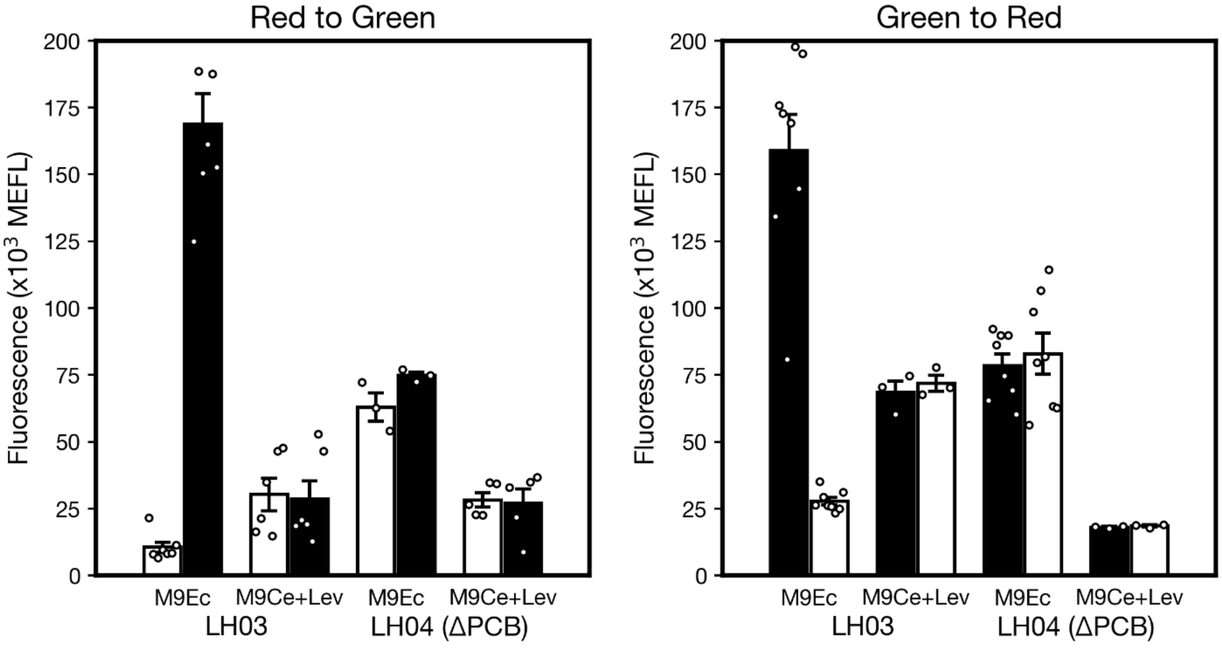


**Supplementary Figure 4. Bacteria on the exterior of worms do not contribute to the measured light response in flow cytometry experiments.** Worms were suspended in clear tubes containing M9Ce+Lev media and levamisole in the flow cytometry experiments. To demonstrate that bacterial cells outside the worm gut (e.g. on the exterior of the worm) do not respond to light in these conditions, samples of pre-conditioned bacteria from the NGM plates were suspended in the M9Ce+Lev or *E. coli* M9 media supplemented with casamino acids and glucose (M9Ec). These samples were then exposed to either green or red light for 8 h, then measured via flow cytometry after the fluorophore maturation protocol (**Methods**). As expected, the bacterial populations are only responsive to light when grown in M9Ec and when the PCB operon is intact (strain JW1935-1/pLH405/pSR43.6, aka LH03). We conclude that the responses shown in data in **Fig. 1e-f** and **Supplementary Fig. 3** are not due to bacteria outside the worm, nor to bacteria that escape the worm over the course of the experiment, as such cells do not respond to light in the experimental buffer (M9Ce). Data represent 7, 6, 3, and 5 replicates (left); and 8, 3, 8, and 3 replicates (right) over 8 and 10 days, respectively; error bars: SEM. Individual data points for each condition are overlaid as white markers.
