## Supplementary Tables for "Optogenetic control of gut bacterial metabolism to promote longevity"

**Supplementary Table 1 | Plasmids used in this study**

| Name | Size (bp) | Resistance | Origin of replication | Description | Reference |
|---|---|---|---|---|---|
| pLH401 | 4,770 | C | ColE1 | Constitutive *ccaR* expression, $P_{cpcG2\text{-}172}$:*sfgfp*, constitutive *mcherry* expression | This study |
| pLH405 | 4,730 | C | ColE1 | Constitutive *ccaR* expression, $P_{cpcG2\text{-}172}$:*gfpmut3**, constitutive *mcherry* expression | This study |
| pLH407 | 2,650 | C | ColE1 | Constitutive *mcherry* expression | This study |
| pLH412 | 6,282 | S | p15A | Constitutive *ccaS(H534A)*, *ho1*, and *pcyA* expression | This study |
| pLH413 | 3,848 | C | ColE1 | Constitutive *ccaR(D51N)* expression, $P_{cpcG2\text{-}172}$:*rcsA* | This study |
| pSR43.6 | 6,282 | S | p15A | Constitutive expression of *ccaS, ho1,* and *pcyA* | Schmidl et al, 2014 |
| pSR49.2 | 4,543 | S | p15A | Constitutive expression of *ccaS* | Schmidl et al. 2014 |
| pMVK201.2 | 3,848 | C | ColE1 | Constitutive expression of *ccaR*, $P_{cpcG2\text{-}172}$:*rcsA* | This study |
| pMVK228 | 4,588 | S | p15A | Constitutive expression of *ccaS* | This study |

C: chloramphenicol, S: spectinomycin

**Supplementary Table 2 | Genbank Accession Numbers**

| Name | Genbank Accession # |
|---|---|
| pLH401 | MN617156 |
| pLH405 | MN617157 |
| pLH407 | MN617158 |
| pLH412 | MN617159 |
| pLH413 | MN617160 |
| pSR43.6 | MN617163 |
| pSR49.2 | MN617164 |
| pMVK201.2 | MN617161 |
| pMVK228 | MN617162 |

### Supplementary Table 3 | Bacterial and worm strains used in this study

| Strain | Plasmids | Background | Resistance | Description | References |
|---|---|---|---|---|---|
| BW25113 | None | BW25113 | None | Keio parent strain | Baba et al., 2006 |
| Δ*rcsA* | None | JW1935-1 | K | Low CA production | Baba et al., 2006 |
| Δ*lon* | None | JW0429-1 | K | High CA production | Baba et al., 2006 |
| LH01 | pLH401,pSR43.6 | JW1935-1 | C,S,K | **Fig. 1,** microscopy | This study |
| LH02 | pLH401, pSR49.2 | JW1935-1 | C,S,K | ΔPCB control for LH01 | This study |
| LH03 | pLH405, pSR43.6 | JW1935-1 | C,S,K | **Fig. 1**, cytometry | This study |
| LH04 | pLH405, pSR49.2 | JW1935-1 | C,S,K | ΔPCB control for LH03 | This study |
| LH05 | pLH407, pSR43.6 | JW1935-1 | C,S,K | mCherry-only control for cytometry | This study |
| LH06 | pMVK201.2,pLH412 | JW1935-1 | C,S,K | CcaS(H534A) control for MVK29 (**Fig. 3**) | This study |
| LH07 | pLH413, pSR43.6 | JW1935-1 | C,S,K | CcaR(D51N) control for MVK29 (**Fig. 3**) | This study |
| MVK29 | pMVK201.2,pSR43.6 | JW1935-1 | C,S,K | Green light-induced CA secretion (**Fig. 3**) | This study |
| MVK46 | pMVK201.2,pMVK228 | JW1935-1 | C,K | ΔPCB control for MVK29 (**Fig. 3**) | This study |

K: kanamycin

| Name | Genotype | CGC Strain ID | Description | References |
|---|---|---|---|---|
| *glo-1* | *glo-1*(zu391) X | JJ1271 | Used for optogenetic induction of GFP expression in the worm gut. Exhibits fewer intestinal granules, which enables lower host background fluorescence. (**Fig. 1**) | Han et al., 2017 |
| *ges-1* | *raxIs51*[$P_{ges\text{-}1}$::*mitoGFP*] | MCW351 | Used for optogenetic induction of CA production in the worm gut. Mito-GFP used to observe mitochondrial morphology. (**Fig. 3**) | Han et al., 2017 |
| *sqt-3* | *sqt-3*(e2117) V | CB4121 | Temperature-sensitive strain used for longevity studies. (**Fig. 4**) | Han et al., 2017 |

**Supplementary Table 4 | Statistical analysis of worm lifespan experiments**

| Replicate #1 | | | | | | | |
|---|---|---|---|---|---|---|---|
| **Strain** | **Light Condition** | **Lifespan** (Mean ± s.e.) | ***p*-value 1** (G vs.R) | ***p*-value 2** (G-H vs. G-L) | ***p*-value 3** ( vs. ΔRcsA) | ***p*-value 3** ( vs. Δlon) | **Total Number** (censor) |
| JW1935-1 | Red | 6.58 ± 0.16 | | | | <0.0001 | 62 (0) |
| | Green-L | 6.50 ± 0.17 | 0.631 | | | <0.0001 | 48 (0) |
| | Green-H | 6.60 ± 0.15 | 0.972 | 0.639 | | <0.0001 | 63 (0) |
| MVK29 | Red | 7.05 ± 0.20 | | | 0.062 | <0.0001 | 64 (0) |
| | Green-L | 8.56 ± 0.23 | <0.0001 | | 0.029 | 0.732 | 66 (0) |
| | Green-H | 9.66 ± 0.28 | <0.0001 | 0.001 | <0.0001 | 0.192 | 59 (0) |
| *Δlon* | Red | 8.72 ± 0.29 | | | <0.0001 | | 46 (0) |
| | Green-L | 8.21 ± 0.31 | 0.368 | | <0.0001 | | 52 (0) |
| | Green-H | 9.06 ± 0.32 | 0.322 | 0.085 | <0.0001 | | 49 (0) |

| Replicate #2 | | | | | | | |
|---|---|---|---|---|---|---|---|
| **Genotype** | **Light Condition** | **Lifespan** (Mean ± s.e.) | ***p*-value 1** (G vs.R) | ***p*-value 2** (G-H vs. G-L) | ***p*-value 3** ( vs. ΔRcsA) | ***p*-value 3** ( vs. Δlon) | **Total Number** (censor) |
| JW1935-1 | Red | 6.64 ± 0.18 | | | | <0.0001 | 45 (0) |
| | Green-L | 6.46 ± 0.16 | 0.481 | | | <0.0001 | 59 (0) |
| | Green-H | 6.61 ± 0.17 | 0.933 | 0.519 | | <0.0001 | 56 (0) |
| MVK29 | Red | 7.00 ± 0.24 | | | 0.157 | <0.0001 | 50 (0) |
| | Green-L | 8.57 ± 0.25 | <0.0001 | | <0.0001 | 0.672 | 56 (0) |
| | Green-H | 9.66 ± 0.27 | <0.0001 | 0.002 | <0.0001 | 0.222 | 58 (0) |
| *Δlon* | Red | 8.73 ± 0.29 | | | <0.0001 | | 48 (0) |
| | Green-L | 8.15 ± 0.33 | 0.291 | | <0.0001 | | 47 (0) |
| | Green-H | 9.08 ± 0.28 | 0.322 | 0.052 | <0.0001 | | 61 (0) |

Censor worms are those that are lost or exhibit body bursting
